# Epidermal Green Autofluorescence is A Novel Biomarker for Local Inflammation of the Skin

**DOI:** 10.1101/2021.02.21.432139

**Authors:** Yujia Li, Weihai Ying

## Abstract

Inflammation of the skin is not only a key pathological factor of multiple major skin diseases, but also a hallmark of the pathological state of the skin. However, there has been significant deficiency in the biomarkers and approaches for non-invasive evaluations of local inflammation of the skin. In our current study, we used a mouse model of skin inflammation to test our hypothesis that the inflammation of the skin can lead to increased epidermal green autofluorescence (AF), which can become a novel biomarker for non-invasive evaluations of the local inflammation of the skin. We found that 12-*O*-tetradecanoylphorbol-13-acetate (TPA), a widely used inducer of skin inflammation, induced not only inflammation of the skin, but also increased green AF of the skin. The distinct polyhedral structure of the increased AF has indicated that the AF originates from the keratin 1 and/or keratin 10 of the suprabasal keratinocytes. Our Western blot assays showed that TPA produced dose-dependent decreases in the levels of both keratin 1 and keratin 10, suggesting that TPA produced the increased epidermal green AF at least partially by inducing cleavage of keratin 1 and/or keratin 10. Collectively, our study has indicated that epidermal green AF is a novel biomarker for non-invasive evaluations of the local inflammation of the skin. This finding is of profound and extensive significance for non-invasive and efficient diagnosis of multiple inflammation-mediated skin diseases. This biomarker may also be used for non-invasive and rapid evaluations of the health state of the skin.

## Introduction

Inflammation is a critical pathological factor in multiple major diseases including cerebral ischemia (1), lung cancer (2) and acute myocardial infarction (3). Local inflammation of the skin is also a crucial common pathological factor in multiple skin diseases, including Psoriasis (4), atopic dermatitis (5), systemic sclerosis (6), and systemic lupus erythematosus (7). In therapeutic aspect, skin’s inflammation is a key therapeutic target for such skin diseases as psoriasis. Therefore, evaluations of the inflammatory state of the skin are crucial for both diagnosis and therapy of the skin diseases.

The current major approach for evaluations of local inflammation of the skin is based on the observations by physicians on pathological changes of the skin such as redness and swollen levels of the skin. These approaches are not quantitative approaches, which also lack sensitivity. Although blood tests can provide quantitative results of the inflammation in the blood, the major limitation of the blood tests is that the tests can provide only the levels of systemic inflammation. Blood tests also have other limitations: The methods are invasive, time consuming and relatively expensive. Therefore, it is of profound clinical significance to search for non-invasive, quantitative, efficient and economic approaches for evaluating the local inflammation of the skin.

Our previous studies have found that LPS, an inducer of systemic inflammation, can produce increased skin’s green autofluorescence (AF) of mice’s ears (8, 9). Moreover, we found that the LPS dosages are significantly associated with the epidermal green AF intensity (8). Based on these observations, we proposed our hypothesis that local inflammation of the skin may also induce increased green AF, which may become a novel biomarker for evaluating the local inflammation of the skin. In our current study, we used a mouse model of TPA-induced skin inflammation to test this hypothesis. Our studies have provided evidence supporting this hypothesis.

## Materials and Methods

### Materials

All chemicals were purchased from Sigma (St. Louis, MO, USA) except where noted.

### Animal Studies

Male C57BL/6BL/6Slac mice of SPF grade were purchased from SLRC Laboratory (Shanghai, China). The animal protocols were approved by the Animal Study Committee of the School of Biomedical Engineering, Shanghai Jiao Tong University.

### Administration of TPA

Male C57BL/6BL/6Slac mice with the weight of 20 - 25 g were administered with 1.0 or 2.5 μg/ear TPA. The stock solution of TPA with the final concentration of 1.0 mg/ml was made by dissolving TPA in acetone. Acetone was used for dilutions of the stock solution of TPA. The mice’s ears were administered with 1.0 or 2.5 μg/ear TPA (20 μl/ear) once. Twenty-four hours after the administration, the skin’s green AF of the mice’s ears were imaged and quantified.

### Imaging of epidermal AF of mice

As described previously (10), the epidermal AF of the ears of the mice was imaged under a fluorescence microscope (A1 plus, Nikon Instech Co., Ltd., Tokyo, Japan), with the excitation wavelength of 488 nm and the emission wavelength of 500 - 530 nm. The AF was quantified as follows: Sixteen spots with the size of approximately 100 × 100 μm^2^ on the scanned images were selected randomly. After the average AF intensities of each layer were calculated, the sum of the average AF intensities of all layers of each spot was calculated, which is defined as the AF intensity of each spot.

### Western blot assays

As described previously (10), the lysates of the skin were centrifuged at 12,000 *g* for 20 min at 4°C. The protein concentrations of the samples were quantified using BCA Protein Assay Kit (Pierce Biotechonology, Rockford, IL, USA). As described previously(11), a total of 40 μg of total protein was electrophoresed through a 10% SDS-polyacrylamide gel, which were then electrotransferred to 0.45 μm nitrocellulose membranes (Millipore, CA, USA). The blots were incubated with a monoclonal Anti-Cytokeratin 1 (ab185628, Abcam, Cambridge, UK) (1:4000 dilution), Anti-Cytokeratin 10 (ab76318, Abcam, Cambridge, UK) (1:1000 dilution) or actin (1:1000, sc-58673, Santa Cruz Biotechnology, Inc., Dallas, TX, USA) with 0.05% BSA overnight at 4°C, then incubated with HRP conjugated Goat Anti-Rabbit IgG (H+L) (1:4000, Jackson ImmunoResearch, PA, USA) or HRP conjugated Goat Anti-mouse IgG (1:2000, HA1006, HuaBio, Zhejiang Province, China). An ECL detection system (Thermo Scientific, Pierce, IL, USA) was used to detect the protein signals. The intensities of the bands were quantified by densitometry using Image J.

### Statistical analyses

All data are presented as mean + SEM. *P* values less than 0.05 were considered statistically significant.

## Results

The ears of C57BL/6BL/6 mice were treated with various dosages of TPA - an inducer of skin inflammation. TPA dose-dependently induced increases in the redness of the ears’ skin (Fig. 1), suggesting that TPA produced increased inflammation of the skin.

**Figure 1.**
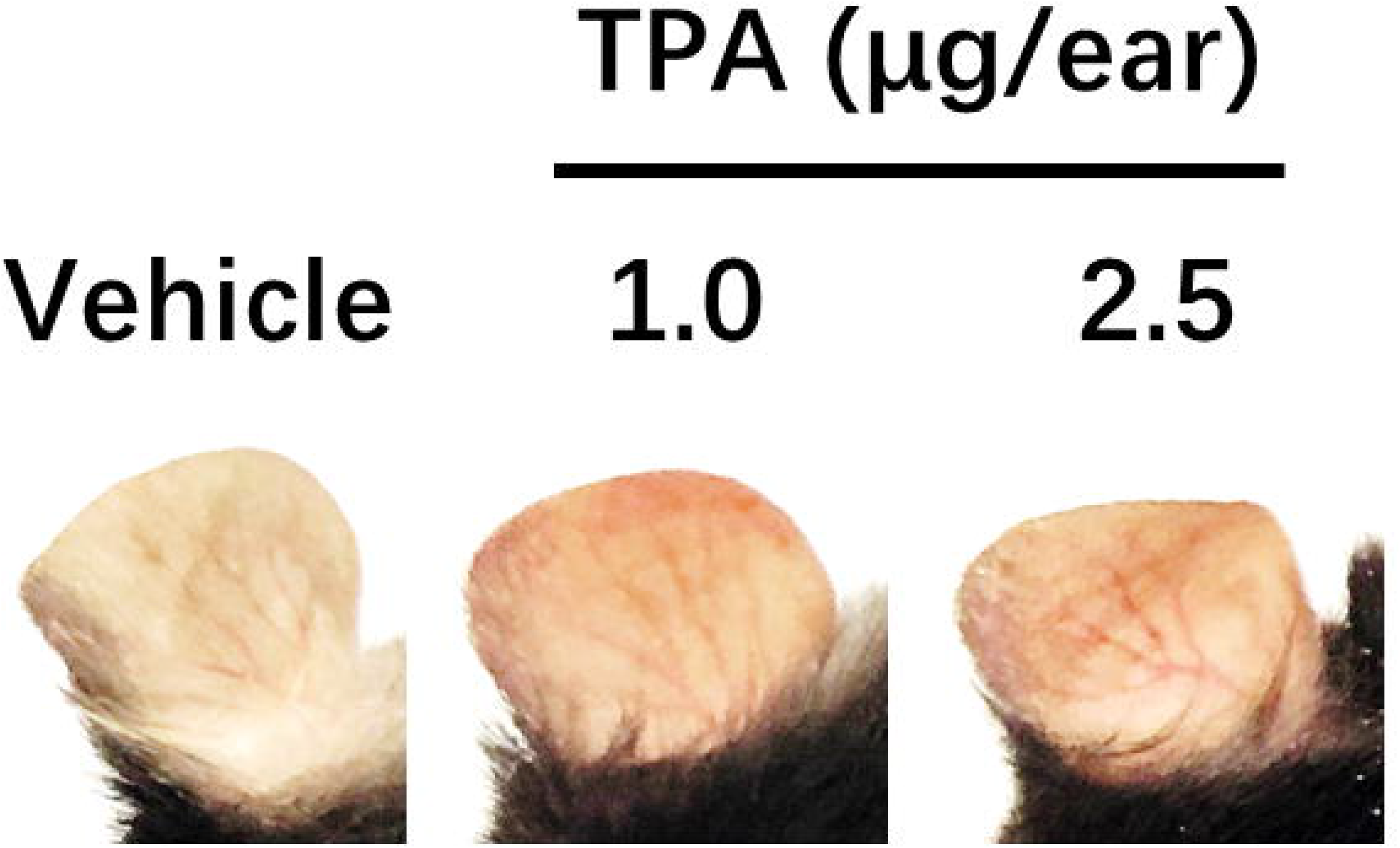
TPA induced redness and swelling of C57BL/6 mouse’s ears. Administration of 1.0 or 2.5 μg/ear TPA induced redness and swelling of the ear’s skin of C57BL/6 mice, photographed at 24 h after the TPA administration. N = 10 - 20.

We determined the green AF of the mice’s ears 24 h after the TPA exposures. TPA produced increases in the green AF of the skin (Fig. 2A). Quantifications of the AF showed that TPA produced significant and dose-dependent increases in the AF (Fig. 2B). The images of the increased AF showed polyhedral structure, which is the characteristic structure of the suprabasal keratinocytes of the spinous layer (12, 13).

**Figure 2.**
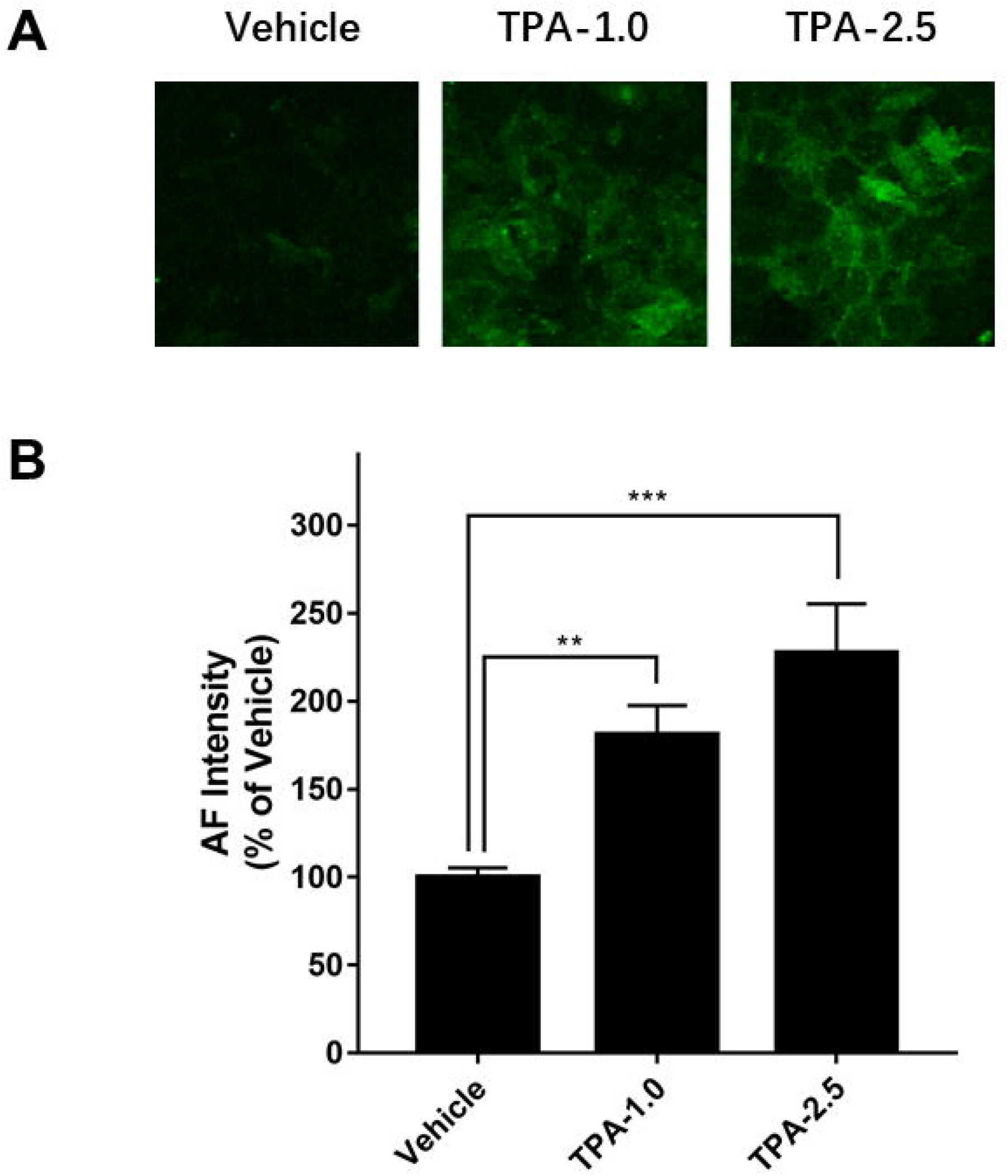
TPA induced dose-dependent increases in the epidermal green autofluorescence of C57BL/6 mouse’s ears. (A) Administration of 1.0 or 2.5 μg/ear TPA increased the skin’s green AF of C57BL/6 mice’s ears, assessed 24 h after the TPA administration. Scale bar = 20 μm. (B) Quantifications of the AF of the images showed that both 1.0 and 2.5 μg/ear TPA induced significant and dose-dependent increases in the skin’s green AF of C57BL/6 mice’s ears. N = 10 - 20. **, *P* < 0.01; ***, *P* < 0.001.

The sole autofluorescent molecules that selectively exist in the suprabasal keratinocytes of the spinous layer are keratin 1 (K1) and keratin 10 (K10) (14). Since our previous studies have suggested that UVC exposures induce increased green AF of the ear’s skin by producing degradation of K1 (15), we determined the levels of K1 and K10 in the ears of the mice 24 h after the mice were administered with TPA: TPA (2.5 μg/ear) produced a significant decrease in K1 (Figs. 3A and 3B). TPA also produced significant and dose-dependent decreases in K10 (Figs. 3A and 3C).

**Figure 3.**
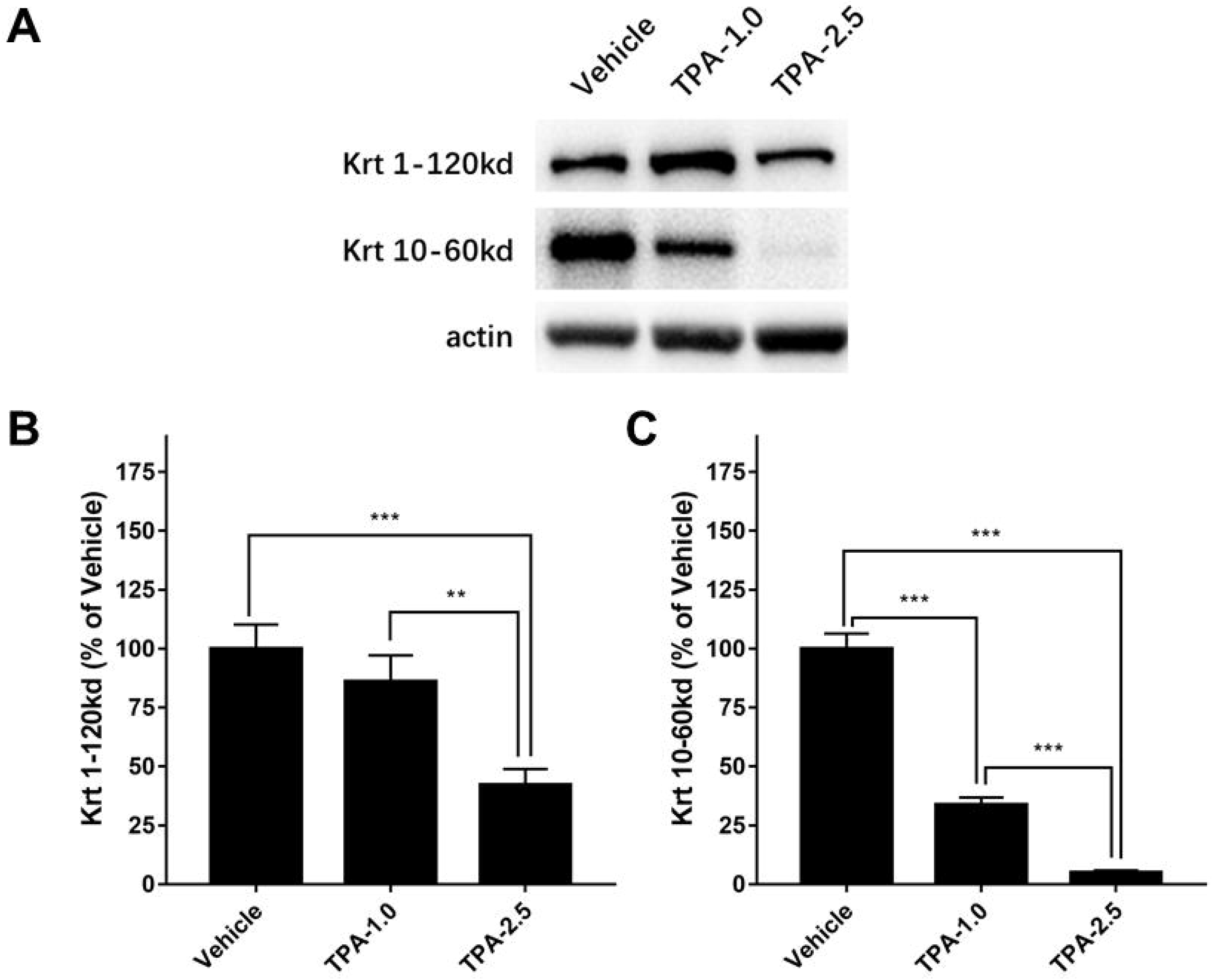
TPA administration produced dose-dependent decreases in the levels of both keratin 1 and keratin 10 of C57BL/6 mouse’s ears. (A) Western blot assays showed that administration of 1.0 or 2.5 μg/ear TPA led to decreased levels of keratin 1 and keratin 10 of the mouse’s ears, assessed 24 h after the TPA administration. (B) Quantifications of the Western blot showed that 2.5 μg/ear TPA produced a significant decrease in the keratin 1 level of the ears. (C) Quantifications of the Western blot showed that both 1 μg/ear and 2.5 μg/ear TPA produced significant decreases in the keratin 10 level of the ears. N = 6. *, *P* < 0.05; **, *P* < 0.01; ***, *P* < 0.001.

## Discussion

The major findings of our current study include: First, local inflammation of the skin can produce significant and dose-dependent increases in the green AF with polyhedral structure of the skin; and second, local inflammation of the skin can also produce dose-dependent and significant decreases in the levels of both keratin 1 and keratin 10, suggesting that the local inflammation-induced cleavage of K1 and/or K10 may mediate the increases in epidermal green AF.

Inflammation is a key common pathological factor of multiple skin diseases. Therefore, evaluations of the inflammation state of the skin are crucial for diagnosis of the diseases. The current major approach for evaluations of skin’s inflammation is observations by physicians on such pathological changes of the skin as redness, which is not a quantitative approach. Determinations of the inflammation levels by blood tests can provide quantitative results, while the blood tests have multiple limitations. Therefore, it is of great clinical significance to search for non-invasive, quantitative, efficient and economic approaches for evaluating the skin’s inflammation.

Our current study has found that TPA produced dose-dependent increases in the green AF of the skin. This finding has suggested a novel, non-invasive, rapid and economic approach for quantitative evaluations of the inflammation levels of the skin. Moreover, when it produced only a mild increase in the redness of the skin, 1 μg/ear TPA produced a significant increase in the green AF intensity, suggesting that our approach is a sensitive approach for evaluating the local inflammation levels of the skin. To our knowledge, our approach is the first non-invasive, quantitative, efficient and economic approach for evaluating the skin’s inflammation. Since it is of crucial importance to evaluate the levels of skin’s inflammation quantitatively and non-invasively, our approach may hold great promise in diagnosis of multiple skin diseases. With increases of the biomedical data on this topic, additions of clinical symptoms, and AI applications, it is expected that our approach will be increasingly sensitive and precise in diagnosis of multiple skin diseases.

The images of the TPA-induced increases in the green AF also showed polyhedral structure, which is the characteristic structure of the suprabasal keratinocytes of the spinous layer of the epidermis (12, 13). The structure of the AF is highly similar to that of the skin of the mice exposed to UVC and LPS (14). Since our previous studies have suggested that UVC exposures induce increased green AF of the ear’s skin by producing degradation of K1 (14), we determined the effects of TPA on the K1 and K10 levels of the ears. We found that TPA produced significant and dose-dependent decreases of both K1 and K10. Collectively, these observations have suggested that TPA could induce increased green AF by producing degradation of K1 and/or K10. It is warranted to further investigate the mechanisms underlying the skin inflammation-induced increases in the green AF.

Since increased skin’s inflammation is an index of pathological state of the skin, our skin’s green AF-based approach may also be used for non-invasive and rapid evaluation of the health state of skin. Due to the importance and extensiveness of applications for evaluating the skin’s health state, it is expected that our approach for evaluating local inflammation of the skin may also have extensive applications in physical examinations and cosmetics industry.

## Acknowledgment

The authors would like to acknowledge the financial support by a Major Special Program Grant of Shanghai Municipality (Grant # 2017SHZDZX01) (to W.Y.) and a Major Research Grant from the Scientific Committee of Shanghai Municipality #16JC1400500 and #16JC1400502 (to W.Y.).

